# Depletion of the *Nb*CORE receptor drastically improves agroinfiltration productivity in older *Nicotiana benthamiana* plants

**DOI:** 10.1101/2023.01.18.517935

**Authors:** Isobel Dodds, Changlong Chen, Pierre Buscaill, Renier A. L. van der Hoorn

## Abstract

*Nicotiana benthamiana* is increasingly used for transient gene expression to produce antibodies, vaccines, and other pharmaceutical proteins but transient gene expression is low in fully developed, 6-8wk old plants. This low gene expression is thought to be caused by the perception the cold shock protein (CSP) of *Agrobacterium tumefaciens*. The CSP receptor is contested because both *Nb*CSPR and *Nb*CORE have been claimed to perceive CSP. Here, we demonstrate that CSP perception is abolished in 6wk-old plants silenced for *Nb*CORE but not *Nb*CSPR. Importantly, older *Nb*CORE-silenced plants support a drastically increased level of GFP fluorescence and protein upon agroinfiltration. The drastic increase in transient protein production in *Nb*CORE depleted plants offers new opportunities for molecular farming, where older plants with larger biomass can now be used for efficient protein expression.

---

*Nicotiana benthamiana* is frequently used for transient gene expression (Bally et al., 2018). In addition to studies on subcellular localization, protein-protein interaction and enzymatic activities, transient gene expression is commercially used to produce antibodies, vaccines, and other pharmaceutical proteins (Sainsbury, 2020; Schillberg & Spiegel, 2022). Transient expression is achieved by infiltrating leaves or whole plants with disarmed *Agrobacterium tumefaciens* harboring a binary vector that carries genes-of-interest on the transfer DNA (T-DNA). *A. tumefaciens* transfers this T-DNA into the plant cell, where it is expressed.

The success of transient gene expression strongly depends on plant age (Saur al., 2016). Best expression upon agroinfiltration is achieved in 3-5 week (wk) old plants. In contrast, poor transient gene expression is achieved in older, 6-8wk old plants that start flowering, despite having more biomass and large leaves that are easy to infiltrate (Lai & Chen, 2013; Saur al., 2016). The poor gene expression is thought to be caused by the perception of cold shock protein (CSP) of *A. tumefaciens* (Saur al., 2016). A 22 amino acid fragment of CSP called csp22 is sufficient to trigger immune responses including a burst of reactive oxygen species (ROS) (Felix & Boller, 2003; Saur et al., 2016). The csp22-induced ROS burst is released from leaf discs from old plants, but not from young plants, implicating that CSP recognition is age dependent and underpins the limitation of transient gene expression in older plants (Saur et al., 2016).

Two receptors have been proposed to perceive csp22 and both are transcriptionally induced in older *N. benthamiana* plants (Saur et al., 2015; Wang et al., 2016). Receptor-like protein *Nb*CSPR was reported to interact with csp22 and required for its perception, because depletion of *Nb*CSPR by virus-induced gene silencing (VIGS) suppresses csp22-induced ROS responses (Saur et al., 2015). In addition, a receptor-like kinase recognizing csp22 has been identified from tomato and is called CORE (Wang et al., 2016). The *N. benthamiana* ortholog of tomato CORE (*Nb*CORE) shares only 29.9% amino acid identity with *Nb*CSPR and binds csp22 with high affinity and specificity (Wang et al., 2016). *Nb*CORE confers csp22 responsiveness when transformed into *Arabidopsis thaliana*, which is otherwise insensitive to csp22 (Wang et al., 2016).

Here, we depleted *Nb*CSPR and *Nb*CORE by VIGS to investigate if CSP perception hampers recombinant protein production in older plants. To silence *Nb*CSPR, we re-synthesized the exact same 299 bp gene fragment of *Nb*CSPR used earlier (Saur et al., 2016; Supplemental **Table S1**), and cloned this into a vector expressing *RNA2* of tobacco rattle virus (TRV2gg). Similarly, a 300 bp fragment specific to *Nb*CORE was cloned into TRV2gg. Alignments of the used silencing fragments with the coding sequences of *Nb*CORE and *Nb*CSPR show that cross-silencing is unlikely (Supplemental **Figure S1**). TRV carrying a fragment of beta-glucuronidase (*TRV::GUS*) was included as a negative control. 2wk-old *N. benthamiana* plants were infected with TRV carrying the silencing fragments. The *TRV::NbCSPR* and *TRV::NbCORE* plants have no developmental phenotypes compared to *TRV::GUS* plants (**Figure 1A**).

**Figure 1.**
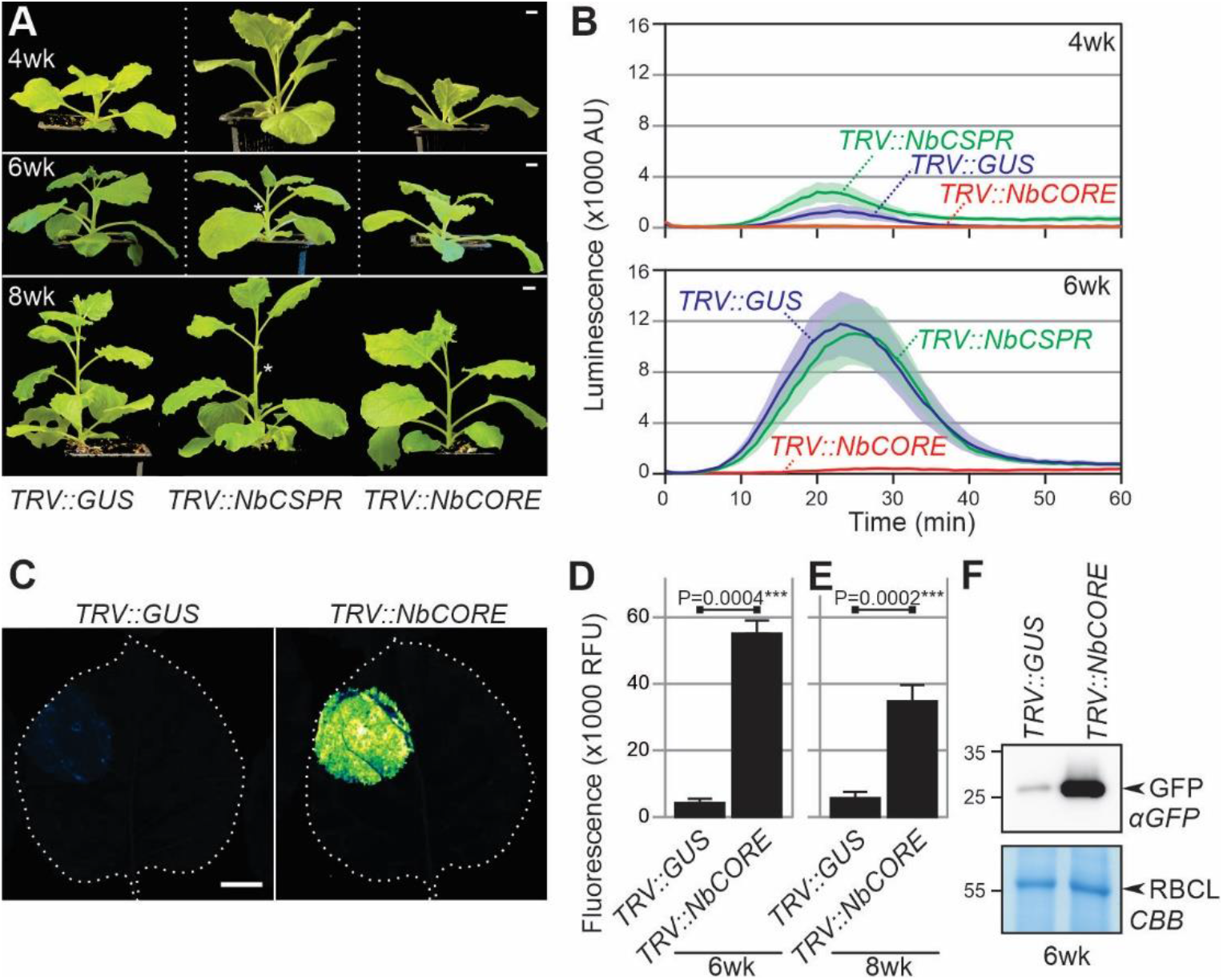
*Nb*CORE silencing removes csp22 responsiveness and increases transient protein production in older *N. benthamiana* plants. **(A)** *TRV::NbCSPR* and *TRV::NbCORE* plants have no additional development phenotype compared to *TRV::GUS* plants. Scale bars, 1 cm. *, removed sample leaves. **(B)** The csp22-induced oxidative burst is absent from 6wk-old *TRV::NbCORE* plants but present in *TRV::GUS* and *TRV::NbCSPR* plants. Error shades represent the standard error of n=6 leaf discs. **(C)** *Nb*CORE depletion causes bright GFP fluorescence upon agroinfiltration of 6wk-old plants. Image was taken five days agroinfiltration with 35S:eGFP. Scale bar, 1 cm. **(D)** Significant increase in GFP fluorescence upon *Nb*CORE depletion. GFP fluorescence was quantified from images of n=4 biological replicates of 6wk-old VIGS plants agroinfiltrated with 35S:eGFP 5 days before fluorescence scanning. Fluorescence was quantified using ImageJ and normalized by leaf area. ****, P value = 0.0000084 (*t*-test). **(E)** *TRV::NbCORE* plants accumulate much more GFP protein upon agroinfiltration than *TRV::GUS* plants. Total leaf proteins were extracted from VIGS plants, 5 days after agroinfiltration with *35S:eGFP*, and analyzed by anti-GFP western blot. CBB, Coomassie brilliant blue.

To investigate which receptor is required for csp22 perception, leaf discs from 4wk and 6wk-old VIGS plants were tested for a csp22-induced oxidative burst. Importantly, the csp22-induced ROS burst is absent from 6wk-old *TRV::NbCORE* plants, and is present in *TRV::NbCSPR* plants, comparable to *TRV::GUS* control plants (**Figure 1B** and Supplemental **Figure S2**). As reported before, younger, 4wk-old plants, only have very weak csp22-induced responses that can vary per batch of plants (**Figure 1B** and Supplemental **Figure S2**). These data demonstrate that *Nb*CORE is essential for the csp22-induced oxidative burst in older *N. benthamiana* plants. Together with the results of Wang et al. (2016), these data show that *Nb*CSPR is neither sufficient nor necessary for csp22 perception.

To investigate to what level the depletion of *Nb*CORE promotes transient gene expression, we agroinfiltrated 6w-old *TRV::GUS* and *TRV::NbCORE* plants with Agrobacterium delivering enhanced GFP driven by a strong 35S promoter (35S:eGFP, Kourelis et al., 2021), and scanned the agroinfiltrated leaves for fluorescence five days later. Bright GFP fluorescence was detected in *TRV::NbCORE* plants, whereas hardly any fluorescence was detected in *TRV::GUS* plants (**Figure 1C**), corresponding to a nearly eight-fold increased GFP fluorescence (**Figure 1D**). A similar increased fluorescence was observed upon infiltrating 8wk-old plants (**Figure 1D**). Western blot analysis confirmed a drastically increased GFP protein level in *TRV::NbCORE* plants compared to the *TRV::GUS* control plants (**Figure 1E**).

Our data showing that *Nb*CORE is required for csp22-induced oxidative burst is consistent with reports that *Nb*CORE binds csp22 with high affinity (Kd = 6 nM, Wang et al., 2016) and that transient expression of *Nb*CORE confers csp22-responsiveness to leaves of young plants (Wei et al., 2018). The csp22-induced ROS burst in *TRV::NbCSPR* plants was similar to the *TRV::GUS* control, which contradicts earlier work (Saur et al., 2016), despite the fact that we used the exact same gene fragment for VIGS. Our data is, however, consistent with experiments that *Nb*CSPR has very low affinity to the csp22 peptide (Kd = 0.3 mM, Nie et al., 2021), and is unable to confer csp22-responsiveness in transgenic Arabidopsis (Wang et al., 2016). Instead, *Nb*CSPR is identical to RE02, the receptor for VmE02, a conserved Cys-rich protein secreted by diverse microbes (Nie et al., 2021).

Our discovery offers new opportunities in molecular farming, where older plants with larger biomass can now be used for efficient transient gene expression. A more durable depletion of *Nb*CORE can be achieved by genome editing, or by engineering *A. tumefaciens* strains to contain a CSP that is no longer recognized by *Nb*CORE. Both approaches will drastically improve transient protein production in older *N. benthamiana* plants, without the need for a license to work with TRV to deplete *Nb*CORE by VIGS.

## Acknowledgements

We thank Tolga Bozkurt for providing *TRV2gg* and *TRV::GUS*; Urzula Pyzio for excellent plant care; Sarah Rodgers and Caroline O’Brian for technical assistance; and Jiorgos Kourelis for useful suggestions and providing 35S::eGFP. This work was supported by grants from the BBSRC Interdisciplinary DTP DDT00060 (ID), Chinese Scholarship Council (CC), BBSRC BB/R017913/1 ‘GH35’ (PB), and ERC-AdG-2020 101019324 ‘ExtraImmune’ (RH).

## Conflict of interest

The authors declare no conflict of interest.

## Author contributions

ID and CC performed experiments and analysed the data; ID, PB and RH designed experiments; ID and RH wrote the paper with help from all authors. All the authors read and approved the final manuscript.

## Supplemental Methods, Figures and Tables Materials & Methods

### Plant cultivation conditions

Wild-type *Nicotiana benthamiana* (LAB) seeds were sown into a 3:1 mix of soil (Sinclair Modular Seed Peat reduced propagation mix) with vermiculite (Sinclair brand Pro Medium) in 7×7 cm square pots and grown at high humidity under transparent plastic covers for 5 days. Seedlings were uncovered and grown first for one week in the greenhouse at 80-120 umol/m^2^/s light and 21°C (night) and 22-23°C (day) in a 16 hour light regime. Plants were then agroinfiltrated with TRV vectors and grown in a growth chamber at 100 µmols/m^2^/sec light and 21°C and 50-60% relative humidity in a 16 hr light regime. Plants were watered three times per week such that no pots were standing in water overnight.

### Construction of plasmids

The fragments in **Table S1** were synthesized by Twist Biosciences and cloned into the golden-gate compatible vector TRV2gg (Duggan et al., 2016) in a BsaI reaction to produce expression plasmids. The fragment sequence published by (Saur et al., 2016) was used to silence *NbCSPR*. The fragment for silencing *NbCORE* was designed using the SolGenomics VIGS tool, SGN-VIGS (Fernandez-Pozo et al., 2015). Plasmids were transformed into *E. coli* DH10β for amplification, were purified and then transformed into *Agrobacterium tumefaciens* GV3101-pMP90. Transformants were selected on plates of LB-agar medium containing 25 μM rifampicin, 10 μM gentamycin and 50 μM kanamycin. A single colony of transformant was cultured in liquid LB containing the same antibiotics.

### Virus-induced gene silencing

Agrobacteria cultures were grown overnight at 28°C in LB medium containing 25 μM rifampicin, 10 μM gentamycin and 50μM kanamycin. The cultures were centrifuged at 3500 x *g* for 10 minutes at room temperature and then resuspended in infiltration buffer (10 mM MES, 10 mM MgCl_2_, 100 μM acetosyringone at pH 5.7). All cultures were diluted to OD_600_=0.5 and a culture carrying a plasmid encoding TRV1 was mixed at a 1:1 ratio with cultures carrying plasmids encoding TRV2 containing fragments to silence *GUS* (negative control), *NbCSPR, NbCORE* and *PDS* (positive control). *Nicotiana benthamiana* plants (LAB) were grown at 21°C under a 16/8 hour light/dark routine in a greenhouse. Both true leaves of 14-day-old plants were agroinfiltrated with the bacterial suspension using a 1 ml syringe without a needle. Three and five weeks later plants were assessed for silencing by checking for bleached leaves in *TRV::PDS* plants.

### ROS assays

The ROS burst assay was performed as described (Buscaill et al., 2019) with the difference that L-012 (Wako Chemical, Japan) was used instead of luminol and the diameter of leaf discs used here was 4 mm rather than 6 mm. Briefly, after incubation in water overnight, one leaf disc (4 mm diameter) was added to 100 µl solution containing 25 ng/µl L-012, 25 ng/µl HRP and 500 nM csp22. Chemiluminescence was measured immediately with the Infinite M200 plate reader (Tecan, Mannedorf, Switzerland) every minute for one hour. The used csp22 peptide is from *Pto*DC3000 (LNGKVKWFNNAKGYGFILEDGK) and was synthesized by Genscript.

### GFP expression and analysis

Agrobacterium GV3101(pMP90) carrying pJK-B2-022 encoding eGFP (Kourelis et al., 2021) were grown overnight at 28°C in LB medium containing 25 μM rifampicin, 10 μM gentamycin and 50 μM kanamycin. The cultures were centrifuged at 3500 x *g* for 10 minutes at 21 °C and then resuspended in infiltration buffer (10 mM MES, 10 mM MgCl_2_, 100 μM acetosyringone at pH 5.7) to a OD_600_=0.5. Expanded leaves of VIGS plants were agroinfiltrated using a needleless syringe. At 5 days post infiltration (dpi) leaves were scanned for fluorescence on a Amersham Typhoon 5 Biomolecular Imager (GE Healthcare Life Sciences, Little Chalfont, UK) using the Cy2 settings. Quantification of fluorescence was performed using ImageJ and normality was tested by a Shapiro-Wilk test and the probability value was calculated with a Well’s t-test. From the same leaves, 1 cm leaf discs were punched, flash-frozen in liquid nitrogen and ground using a pestle. The leaf tissue powder was mixed 3:1 with phosphate-buffered saline and centrifuged for 10 minutes at 13,000 x *g* at 4°C. The total soluble protein supernatant was mixed 1:3 in 4x gel loading buffer (200 mm Tris-HCl (pH 6.8), 400 mm DTT, 8% SDS, 0.4% bromophenol blue, 40% glycerol) and heated at 95°C for 5 minutes.

### Western blot analysis

Proteins were separated on a 12% w/v SDS-PAGE gel, transferred onto a PVDF membrane using the TransBlot Turbo system (Bio-rad, Hercules, CA) and blocked for 1 hour at room temperature in 5% w/v skimmed milk in PBS with 0.01% v/v Tween-20. eGFP was detected using an α-GFP-HRP antibody (Abcam ab6663, 1/5000) also in 5% w/v skimmed milk in PBS with 0.01% v/v Tween-20. Chemiluminescent signals were detected using the SuperSignal™ West Femto Maximum Sensitivity Substrate (Thermo Fisher Scientific, Waltham, MA, USA). eGFP-antibody complex signals were captured using the ImageQuant LAS 4000 (GE Healthcare, Healthcare Life Sciences, Little Chalfont, UK). A corresponding 12% w/v SDS-PAGE gel was run simultaneously and incubated in InstantBlue® Coomassie Protein Stain (Abcam, Cambridge, UK) and then imaged using the Epson Perfection V600 Photo scanner (Epson, Nagano, Japan).

---

**Figure S1.**
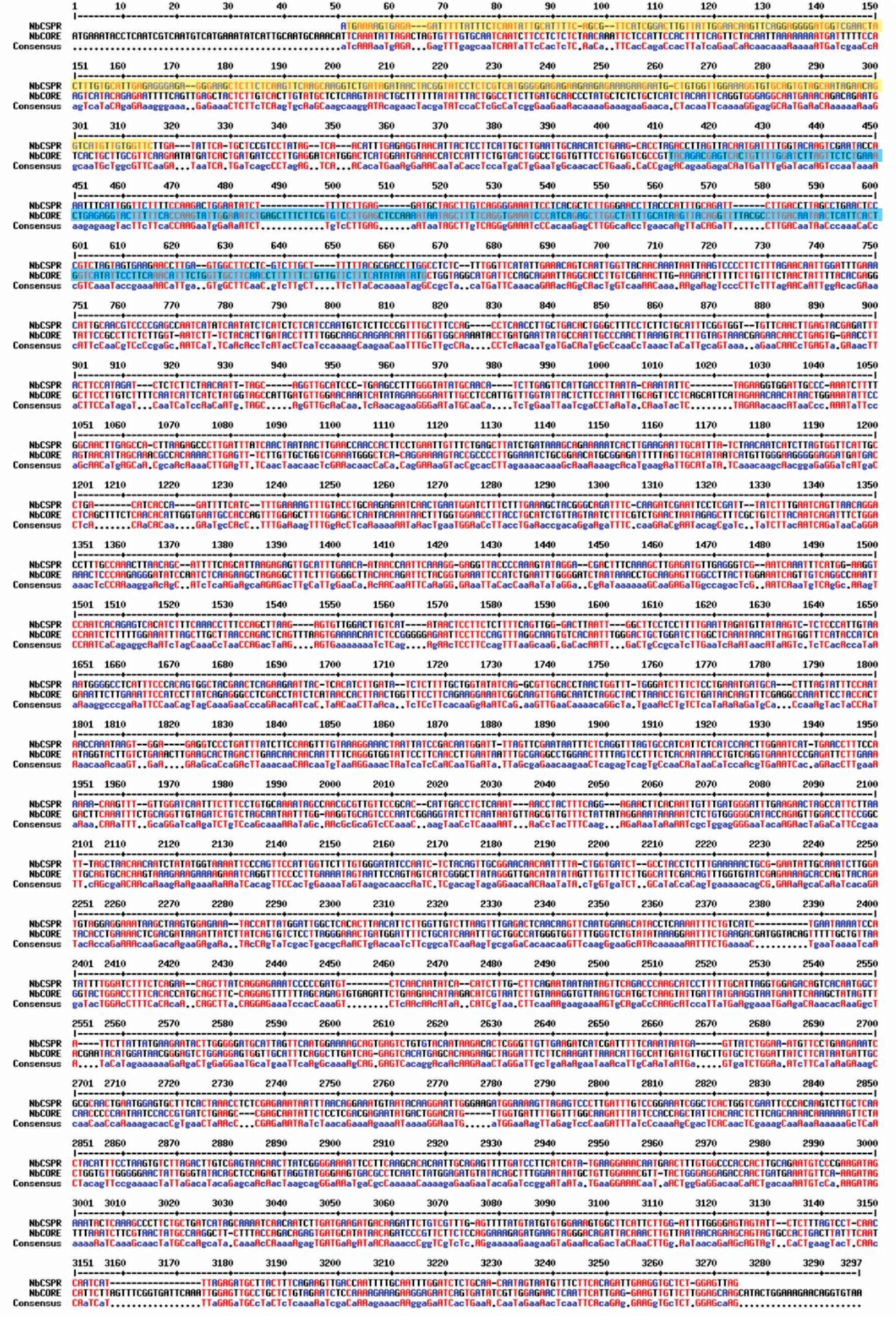
Nucleotide alignment between *Nb*CSPR and *Nb*CORE Nucleotide sequences of the open reading frames of *Nb*CSPR (NbD023129) and *Nb*CORE (NbD017538) were aligned using MultAlin. Fragments used for silencing are highlighted.

**Figure S2.**
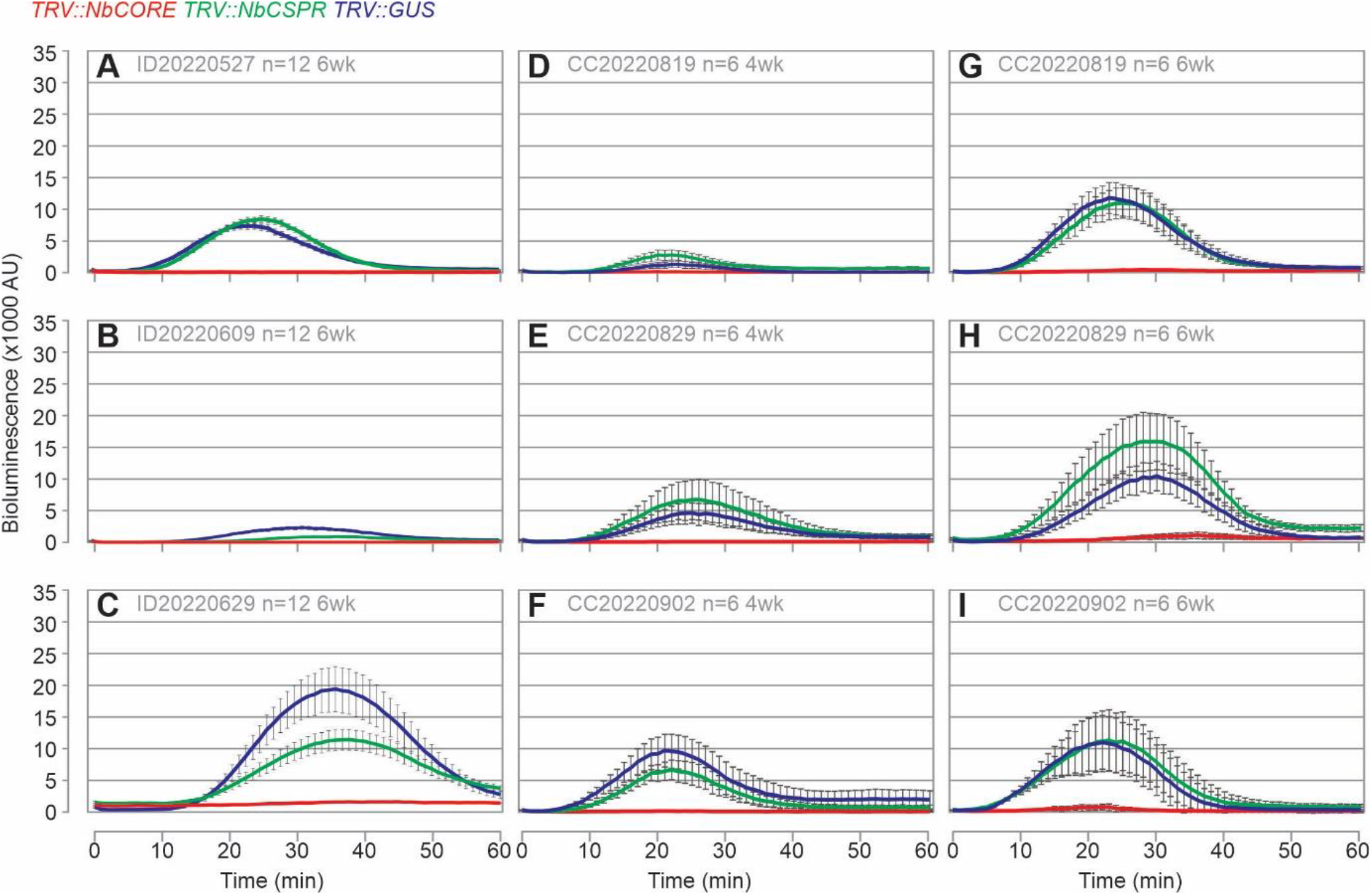
Replicate ROS experiments. Two week (2wk) old plants were agroinfiltrated with a 1:1 mixture of agrobacterium cultures delivering TRV1 and TRV2, respectively. TRV2 contains fragments GUS (blue), *Nb*CSPR (green) or *Nb*CORE (red). Leaf disks were taken from 4wk (D-F) or 6wk (A-C, G-I) old plants, incubated overnight in water and transferred to a 96-well plate. ROS assay solution containing L-012, HRP and 500 nM csp22 peptide of *Pto*DC3000 was added and the luminescence was measured every minute for 60 minutes immediately. Error bars represent standard error of n=12 (A-C) and n=6 (D-I) replicate leaf disks from 3-4 plants. Datasets D and G are also shown as main figure.

**Table S1.**
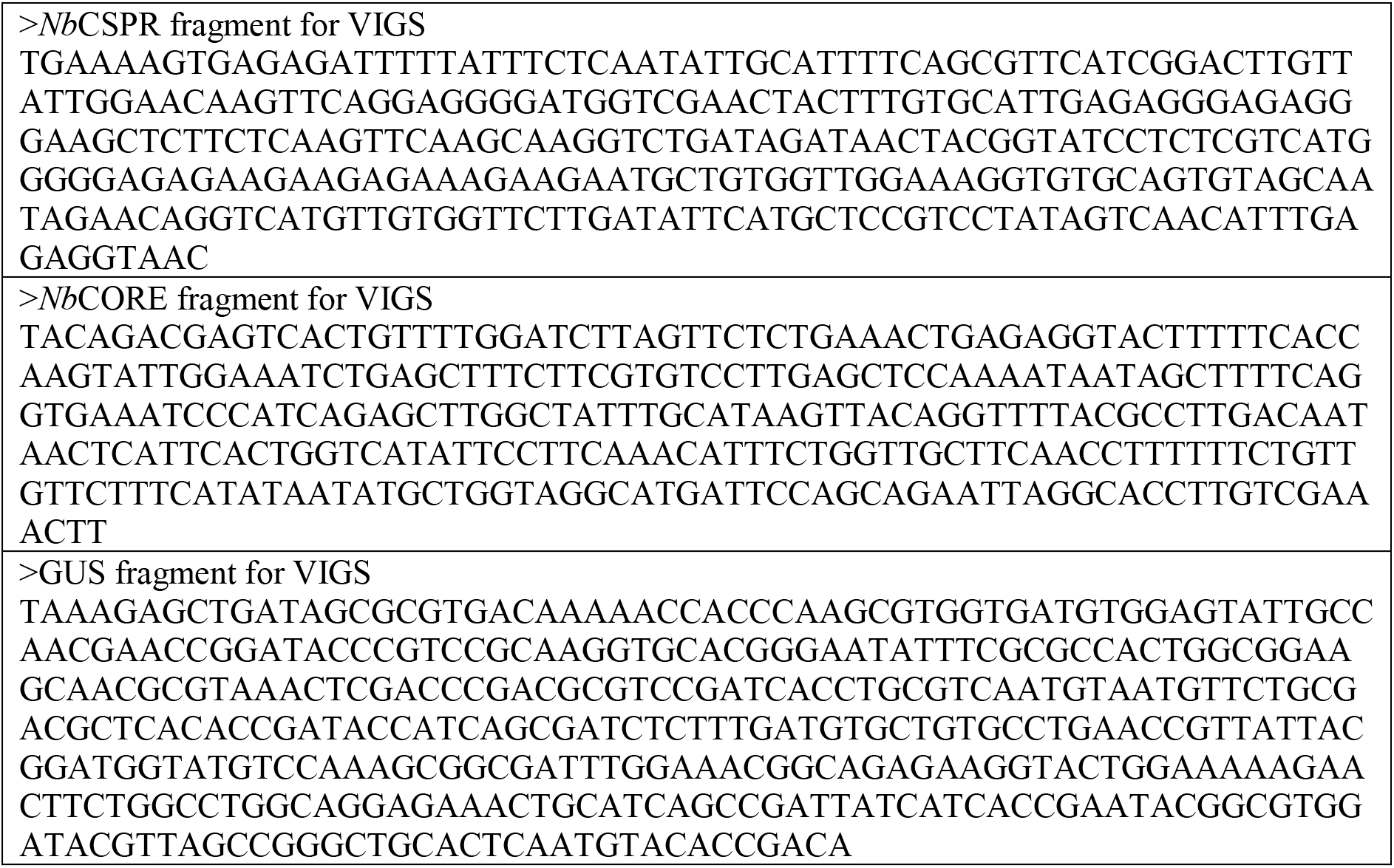
Used oligonucleotides

**Table S2.** Constructs used

| Name | Description | Reference |
| --- | --- | --- |
| pID010 | *TRV2gg::NbCSPR* | This work |
| pID024 | *TRV2gg::NbCORE* | This work |
| TRV::GUS | *TRV2gg::GUS* | Duggan et al., 2021 |
| TRV1 | *RNA1 of TRV* | Liu et al., 2002 |
| pJK-B2-022 | *35S::eGFP* | Kourelis et al., 2021 |
| TRV2::PDS | *TRV2::PDS* | Liu et al., 2002 |

## Notes

### Competing Interest Statement

The authors have declared no competing interest.

